# The kinematics of cyclic human movement

**DOI:** 10.1101/826313

**Authors:** Manfred M. Vieten, Christian Weich

## Abstract

Models describing cyclic movement can roughly be divided into the categories theory or data driven. Theory driven models include anatomical and physiological aspects. They are principally suitable for answering questions about the reasons for movement characteristics. But, they are complicated and substantial simplifications do not allow generally valid results. Data driven models allow answering specific questions but lack the understanding of the general movement characteristic. With this paper we try a compromise not having to rely on anatomy, neurology and muscle function. We hypothesize a general kinematic description of cyclic human motion is possible without having to specify the movement generating processes, and still getting the kinematic right. The model proposed consisting of a superposition of six contributions – subject’s attractor, morphing, short time fluctuation, transient effect, control mechanism and sensor noise -, with characterizing numbers and random contributions. We test the model with data form treadmill running and stationary biking. Applying the model in form of a simulation results in good agreement between measured data and simulation values.

## Introduction

Bipedal gait, especially walking, has been the most decisive development of homo sapiens to surpass its ancestors and relatives [1]. In the past centuries further cyclic motions like swimming, cycling, rowing or skiing came along, to overcome natural obstacles, to facilitate traveling and then as leisure activities. Recently, cyclic motion descriptions served as biological templates for developments in robotics together with developments in artificial intelligence [2]. Although cyclic movements are performed thousand-fold each day in everyday life, their underlying composition and structure is not fully understood. The kinematic of human cyclic motion seems rather simple at first glance. Detailed observations display a repeating structure and some fluctuation producing similar but not identical repetitive cycles of movements [3, 4]. These changes often describe a transient effect at the start of the movement [5, 6, 3], as generally observed in dynamical systems [7, 8]. Moreover, various perturbations alter the regularity of the ongoing movement and stride time dynamics [9-12]. Dingwell and Kang [13] describe these findings as ‘inherent biological noise’, being local instabilities [14] during movements like walking, without causing falls or stumbles, meaning that the subjects move ‘orbitally stable’. Nashner [15] pointed out, that the described continuity after perturbations is retained by adjusting parameters of the present walking motion rather than recruiting a new motor pattern (p. 650). The consequential questions arise after the mathematical structure of the human cyclic movements and how such a developed model can be tested? No wonder that the modern quantitative scientific endeavor to understand the mechanism behind the central movement trait began already as early as the nineteenth century [16]. Describing cyclic motion most often is realized by selected specific body marker and their coordinate portrayal as function of time [17]. The classical gait parameter like step length, step frequency, velocity as well as marker world lines carry most of the information considered. With the advent of direct acceleration measurement further and subtler information, which coordinate explanation cannot deliver, can serve describing cyclic motion. Coordinate data, however, can at least in principle, be generated from acceleration data by two consecutive integrations with respect to time. But, integration is a smoothing process, which makes it evident, that important information gets lost. For this reason, we introduce a *mathematical model of the kinematic of the human cyclic motion* based on acceleration data. It allows simulating cyclic movement and the comparison with measured data. We expound this model as a superposition of six mathematical terms covering the motion as a limit-cycle attractor (1), individual attractor morphing (2), short time random fluctuation in form of “random walk” (3), the transient effect describing initial oscillations around the attractor at the onset of the activity subsiding with increasing time (4), a control process being activated when stride variations tend to exceed the morphed attractors’ boundaries (5), and the influence of noise generated by the measurement device - accelerometers - (6). Thus, this model does allow to extend earlier findings specifically about the variability of subjects’ cyclic movement with its fix and random fractions. There exist two types of models - theory driven and data driven – both with its own strong and weak aspects. For example, a theory driven model as described in [18] gives inside into the working of seven muscle groups within the lower extremities. The necessity of keeping the model manageable, in the mentioned paper by using a 2-dimensional rigid body model, leads to deviations from the real movement. On the other hand the data driven model of [19] was able to detect the influence of emotions onto the movement pattern. They used artificial neural nets, allowing to identify subtle effects. While here the detection movement characteristics caused by emotions is nicely achieved, the specifics of the gait changes remained undetected. With the present paper we attempt a compromise, not having to rely on anatomy and muscle function, but still trying to understand kinematic processes and the movement pattern quantitatively. A study on cycling at two different power outputs (150 W and 300 W) at a cadence of 90 rpm [20] found differences in the muscle activities detected via EMG, while kinematic data stayed almost unchanged. This result together with the stability of the individual’s attractor over time and after rehabilitation [21, 22] is motivation to examine the possibility to quantitatively describe movement without the knowledge of muscle activity. The purpose of this paper is to precisely outline the kinematics of cyclic motion by establishing the necessary mathematical equations, which allow simulation. The method presented permits finding subject specific movement constants. The testing of model and method is done on two classical cyclic motions: running (on a treadmill) and (stationary) biking.

## Method

### Model

We construct the full acceleration as a superposition of the six terms

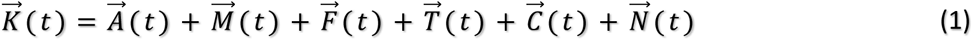

1. 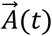 the Limit-Circle-**A**ttractor, a constant acceleration pattern being repeated with every cycle.
2. 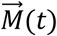 attractor **M**orphing, which allows minor deviations from the actual attractor values.
3. 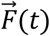 short time **F**luctuation in form of a “random walk”.
4. 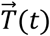 **T**ransient effect, which can be present at start and decreases rapidly.
5. 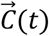 **C**ontrol mechanism, kicking in when actual accelerations deviate too much from the morphed attractor.
6. 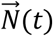 **N**oise caused by the accelerometers.

1. Limit-Circle-Attractor 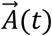 can be regarded as the average of all cycles. This however, is an idealized definition, which cannot fully be met, since this would call for averaging of an infinite number of cycles. Instead, we approximate the attractor by a finite number of cycles, which for later examples we chose as the number of complete cycles within a specified minute of the data collection.

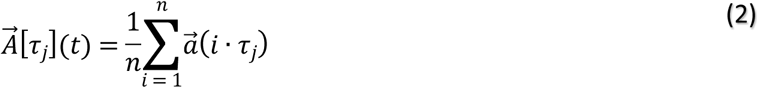

is a closed line in 3D acceleration space with *j* being the number of the data points within an attractor. Such an approximated attractor is characteristic for each individual [21, 22]. The actual calculation starts with dividing each data set into one-minute sections and calculation of the attractors [23]. There is one important methodological difference however. Instead of adding the cycles having different numbers of data points in temporal order, we describe each circle as consisting of a fixed data point number n. This is achieved through spline approximation. n stands for the mean number of data points of all complete cycles within a one-minute interval. Afterwards we add up all cycle values for each of the n points and divide them by the number of cycles. The results represent mean values of the one-minute data sets having the original sampling frequency, still containing the influence of morphing, random walk, transient effect and the control mechanism. A data set least influenced serves as attractor to compare all others with. Appropriate attractors are those for which time *t* ≫ *t* _*T*_ (transient time, explained below).
2. A time-dependent individual attractor morphing 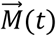 is described as the attractor change from start *t*_*S*_ to end *t*_*E*_ minute by minute. We take care of this process by taking attractor approximations at beginning and end and describe the morphing of the two attractor approximations by

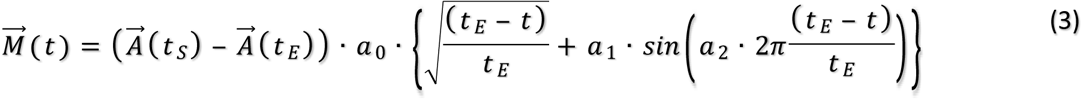 <u>Important to mention:</u> The morphing is small compared to attractor differences between individuals.
3. Fluctuation 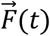 in the form of a “random walk”. These are changes around a morphed attractor described with the iteration

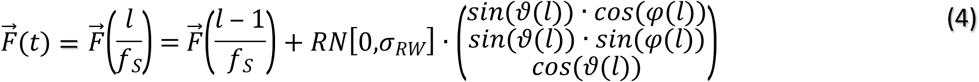 Here, *l* is the data number. An aberration from the attractor can happen in any direction. We describe this using the angles *ϑ* and *φ*. Their actual values are random having a uniform distribution on the sphere with the polar and azimuthal angles:

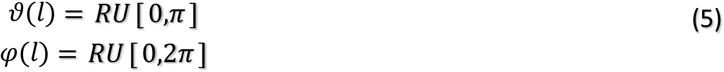 *RU* [*α,β*] represents random generation with a uniform characteristic within the interval [*α,β*]. With this definition the standard deviation of the random walk depends on the sampling frequency *f*_*S*_. Since the random walk must not be depending on the specify of a measurement – the sampling frequency *f*_*S*_ -, we introduce a parameter *ϕ*, which does not change with the sampling frequency.

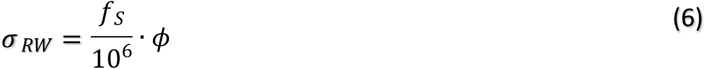 The factor 10^6^ was introduced for convenience. For simulating the movement *ϕ* together with 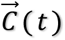 (see below) must be chosen to reproduce the statistical spread of the data around the attractor.
4. The **C**ontrolling mechanism 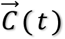, respectively the vector component *C*_*k*_(*t*), is kicking in when the distance to the morphed attractor’s coordinates passes the boarder *b*_*k*_

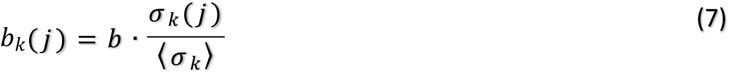

at attractor point *j*. Here *b* is a constant and *σ* _*k*_ (*j*) the attractor’s standard deviation, which is divided by the average of the attractor’s deviation. This takes care of the changing width of the acceleration bundle. The correction term, being activated at time *t*_*b*_, is modeled as

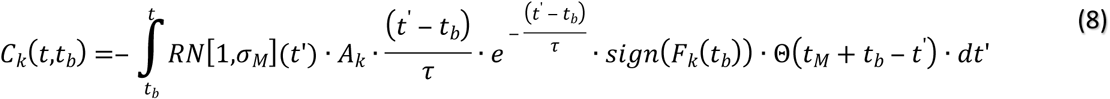 With *sign*(…) being the signum and Θ(…) the step function. We set the maximal acceleration change to *τ* = 80 *ms* in analogy to the style of a muscle’s timely response [24] with acceleration effectively lasting *t*_*M*_ = 4 · *τ* = 320 *ms*, to obtain

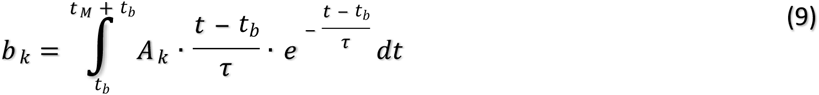

*b* _*k*_ is the acceleration necessary to get back precisely onto the morphed attractor values. This holds true for

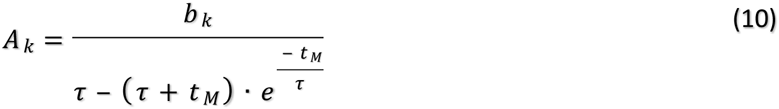 *RN*[1,*σ*_*M*_](*t*) represents a normal distributed random element introducing some deviation from a perfect working controlling mechanism.
5. The transient effect 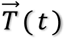 is a temporary oscillation around the attractor at the beginning of a cyclic movement. The starting value of the oscillation might be very individual, specific to the subject, and having a part of the starting value occurring by sheer chance. We model the deviation as the solution of a damped harmonic oscillator, where the transient term can be viewed as the departure from the morphed attractor

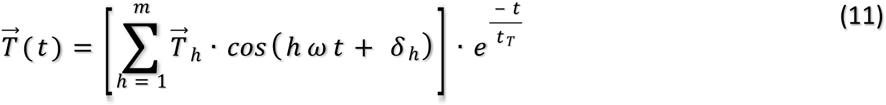 With 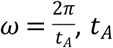 being the average time of one cycle within the one-minute interval Δ*t. δ* _*h*_ is the phase, which within a simulation is chosen randomly being any number between zero and 2*π. h* specifies the number of harmonics contributing, with *m* being the highest one. The maximal harmonic is identified from the Fourier transform of a subject’s movement. *t*_*T*_ denotes the time for the transient effect decreasing to *e* ^− 1^. The transient’s effects value averaged over the n^th^ minute is

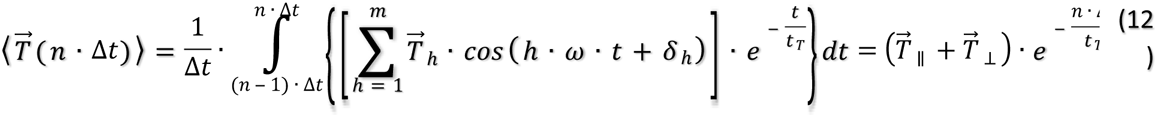 Here and below ∥ stands for the part of the vector pointing in the direction of the combined vectors of 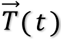 and 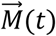. ⊥ indicates the vector parts perpendicular to the mutual direction.
6. Noise (generated by the sensors) 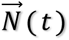 need to be considered. When simulating the kinematics and comparing it with real live data, we need to include the measurement error – noise-caused by the sensor characteristics. It can be obtained directly from measuring the output signals of the sensors at rest. The signal of an accelerometer is, subtracting the values caused by the earth’s gravitational field, modeled as white noise.

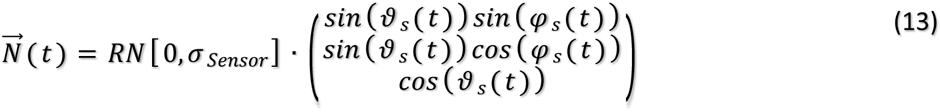

Here *RN* stands for a random normal distributed contribution with a mean value of 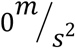 and a standard deviation *σ* _*Sensor*_, which is the characteristic of the specific sensor. *ϑ*_*s*_ and *φ*_*s*_ are randomly chosen to get a uniform distribution on the unique sphere. *σ* _*Sensor*_ is calculated using equation (13) and taking 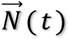 from the data recording of the sensors at rest.

The main parameter for checking the model’s validity is *δM*. It is the average distance between two data sets [23] and is calculated using equation (1) by

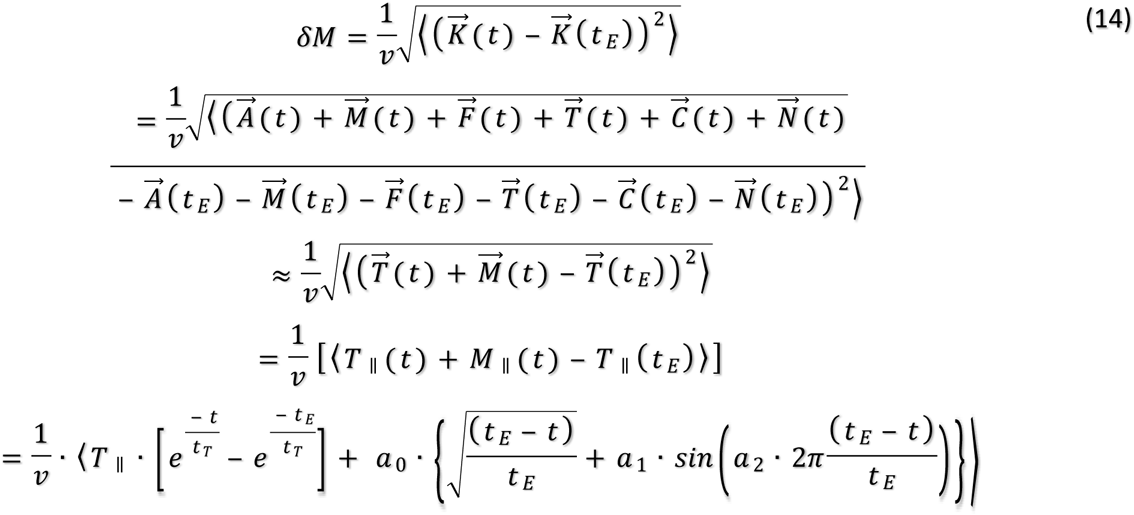

Here 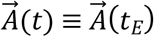 by definition of an attractor as being identical at any cycle. The fluctuation as well as the correction term do have almost identical averaged contributions at the different time intervals of one minute and the noise has contributions almost completely cancelling out within one minute. Therefore, the remaining input comes from the transient effect and the attractor morphing. We calculate the length of the three lasting vectors. The remaining terms are the parallel contributions, all lying in the same direction at a given time, which can be written as a sum of scalars. The subsequent equation allows to write *δM* depending on 5 constants 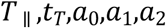, which are specified by curve fitting of the measurements. We use the software CurveExpert Professional 2.6.5, which uses the Levenberg-Marquardt algorithm providing the non-linear curve-fitting. While the three constants on the right describe the highly individual subject and task dependent morphing, the two constants on the left approximate the transient oscillation contributing to *δM* at the beginning of a cyclic movement. *t*_*T*_ depicts the time till the oscillation decreases to 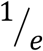 of its original value *T*. The oscillation is neglectable if *t*_*T*_ ≥ *t*_*E*_ (measuring time) since than the two exponential functions are almost identical 1 resulting in these terms to cancel. The values of the morphing and the transient effect do mix, which does not allow to separate these two effects in all cases. Fortunately, there is a method to separate these two effects, which will be explained below. Altogether we have the nine constants 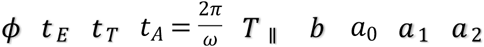 determining our model. All definitions and the respective calculations/approximations are given to allow simulating cyclic motion with the help of the attractors and the constants gained form the measured data. These simulations are naturally not identical to the original data, since the algorithm contains contributions of random processes.

### Subjects

A total of ten athletes, six female and four males, were tested in summer 2019. The running data (n = 5) were collected in Kreuzlingen, Switzerland (Nationale Elitesportschule Thurgau) whereas the cycling measurements (n = 5) took place at the University of Konstanz, Germany. All runners were active experienced leisure athletes. None had suffered any present injury, which could have possibly impeded their performance. The bicyclers were recruited from the local pool of university students. The only prerequisites were to be aged 18 years or older and able to run 60 minutes without reducing their initial pace or cycling at a moderate wattage over 60 minutes regulated by their age, weight and training level [25], respectively. All participants were requested to fill out and sign an informed consent.

### Equipment

To collect the necessary raw accelerometer data, two inertial sensors were used (RehaWatch by Hasomed, Magdeburg, Germany), which were attached to both ankles by a hook-and-loop fastener during the runs and on the proximal frontal part of the tibia (facies medialis) during the cycling tests. The sensors, MEMS - micro-electro-mechanical-system, have a size of 60×35×15 mm and weigh 35 g each. They function as a triaxial accelerometer, which we set up to a measurement interval of ± 8 *g* and triaxial Gyroscope with up to 2000°/*s*. The sampling rate was set to 500 Hz. They measured the acceleration of the feet in three dimensions (x, y, z) with data saved to a smartphone (Samsung Galaxy J5) using the app RehaGait Version 1.3.9 programmed by Hasomed (Magdeburg, Germany). All runs were performed on a treadmill (9500HR by Life Fitness, Unterschleißheim, Germany). The cycling measurements were undertaken on a cycle ergometer (ergoselect200, Ergoline, Bitz, Germany).

### Running data

The first session started with a short 5-minute warm up phase to get familiar with the treadmill and to determine an easy running pace associated with a BORG-scale of 3 [26] (Table 1).

**Table 1.** Running speed of the subjects

|  | Subject 1 | Subject 2 | Subject 3 | Subject 4 | Subject 5 |
| --- | --- | --- | --- | --- | --- |
| Running speed | 10 km/h | 11 km/h | 10 km/h | 8.5 km/h | 8.7 km/h |

The chosen running speed remained stable throughout all following test sessions each lasting 60 minutes. The participating athletes repeated the testing protocol in a time frame of approximately four weeks consisting of five testing days separated by at least 24 hours. The measurements were received from tri-axial accelerometers by a smartphone placed on a desk beside the treadmill to ensure undisturbed reception. Before the actual run the participants set up the treadmill at 1% inclination (to simulate wind resistance) and their individual speed waiting on the collateral standing area close to the treadmill belt. Once the chosen speed of the belt was reached the tester counted down from three to one before starting the data collection on the smartphone. At the same time the runner jumped inwards and started immediately with running at the chosen pace over 60 minutes. This jumping movement, lasting approximately one second, was cut out of the data during the data management process, as it was a running-unspecific movement.

### Cycling data

Within four weeks all cyclists repeated the testing protocol five times. Before the initial test day, the research group calculated the power and selected a convenient seat position. All participants were tested at their preferred cadence (rpm = repetitions per minute), which the participants were able to hold constant over 60 minutes. Their power output conformed with an easy endurance workout and was defined using the athletes’ age, weight and training level [25] (Table 2).

**Table 2.**
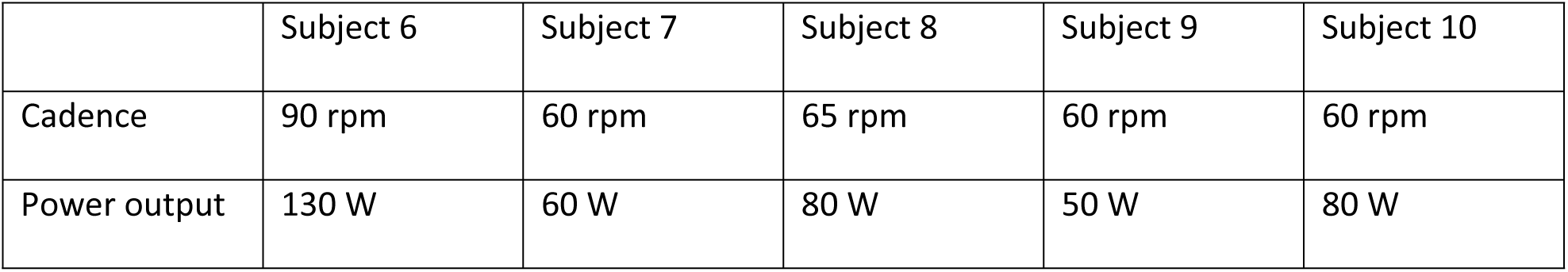
Power output of subjects in biking

On each test day, the cyclists adjusted the seat and the handlebars as determined. The research assistant advised the athlete to hold the seating position and the cadence as stable as possible. The data collection was started by the tester right after the start signal caused the participant to pedal.

### Simulation

For the simulation we created an app called “TrackSimulator”, available as Windows and macOS versions. It was created within Matlab and being available as stand-alone solution without the need to install the Matlab program. The app included all the algorithms described above. As input serve attractors of the tested subjects and the nine, also individuum specific, constants of the model. We set the number of harmonics = 2 within equation (11), because those harmonics carry the majority of the signal’s strength.

### Data handling

Since further analysis required the collected 60-minute data block to be divided into 60-second intervals, a file splitter was applied to produce suitable single datasets. A raw data text-file contained thirteen columns: time and the acceleration as well as the gyro meter data in x, y and z direction for the left and the right foot, respectively. Afterwards an app called “Attractor”, programmed with Matlab was used to calculate the attractor data of every one-minute data set. Each attractor dataset contained 25 x n velocity/cadence-normalized data points: t, x_left foot,_ y_left foot,_ z_left foot,_ x_right foot,_ y_right foot_, z_right foot_, their standards deviations, standard errors, and gyroscope data. The functionality of the Attractor App is based on the attractor method developed by Vieten et al. [23] with the alteration of the attractor building process described above. The attractors were normalized for velocity in running and cadence (normalization factor v=cadence/10) for biking.

### Super attractor

For each participant the final 50 minutes of each run were gathered in a 50 × 25 × n matrix (minute x attractor time/acceleration/deviation/standard error/gyroscope data x average number of attractor points). The first ten minutes were excluded to eliminate the influence of the transient effect. Before carrying out the final analysis, the average of the five matrices were calculated. The resultant attractor is termed *super attractor*.

### Similarity analysis

For this procedure each attractor is recalculated having 500 data points by adjusting the sampling frequency using spline approximation. To find out how similar two movements are, we calculated the *recognition horizon* around each single attractor point, which is defined as the surface area at a distance equal to five standard deviations away from the attractor point. A test attractor is checked point to point if lying in- or outside the *recognition horizon* of the first attractor using another Matlab procedure (Fig 1).

**Fig 1.**
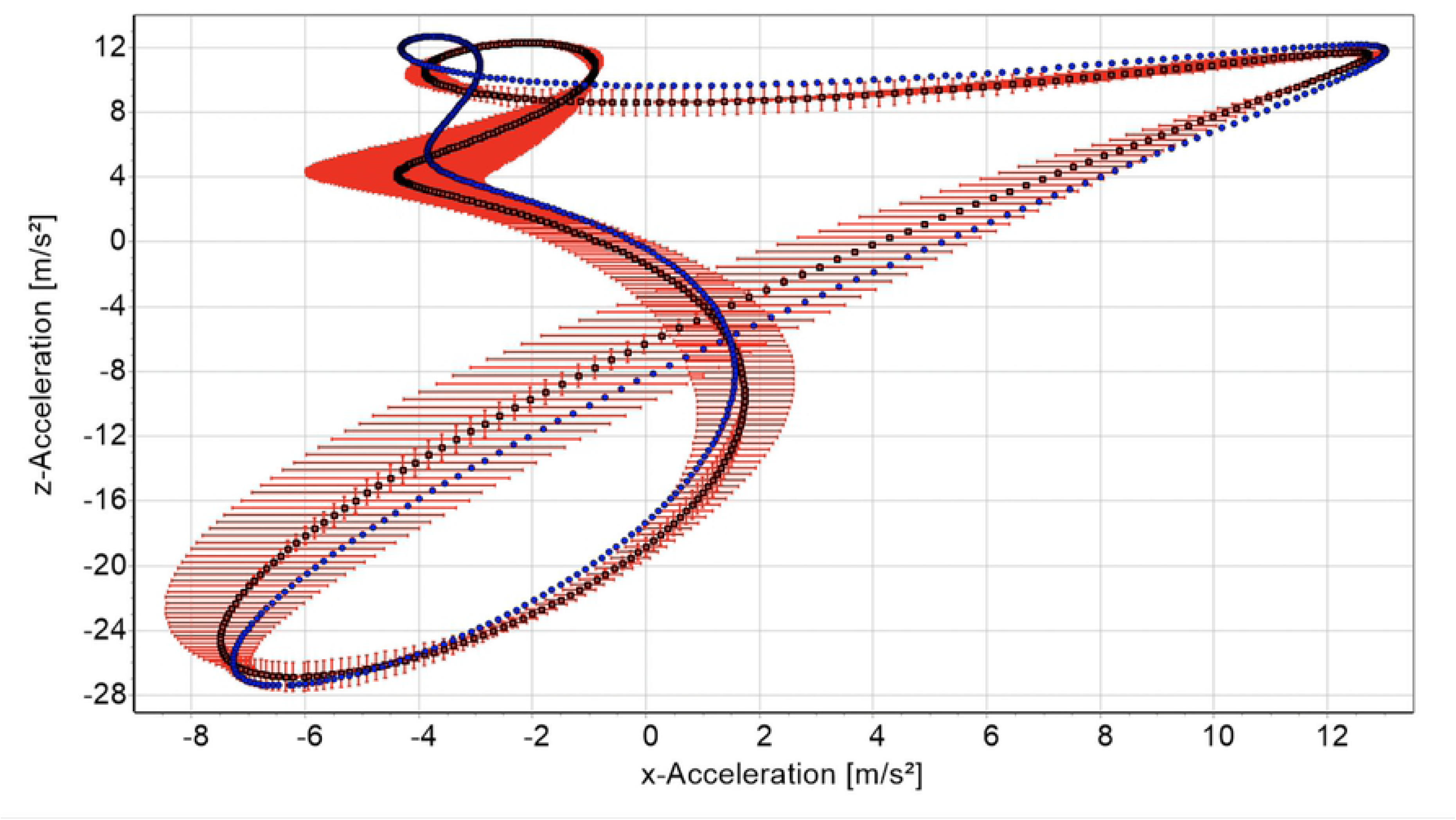
Schematic two-dimensional depiction of the three-dimensional recognition horizon (red) and compared attractor (blue). Each measured or simulated minute over all running or cycling sessions (5 x 60 minutes) got checked against the respective *super attractor*. The *similarity rate* is defined as percentage of data points lying within the *recognition horizons*.

### Separating the transient effect from morphing

To exclude the influence of the morphing as far as possible, we calculated a *super attractor* from 5 independent 1-hour-runs of each individual taken about 5 months before the actual measurements for running. For biking we did not have the data from months before. Here a *super attractor* was created out of four datasets to compare with the fifth one. Since our hypothesis was that an attractor is stable only within a given interval, the *super attractor* represents just one possible attractor configuration. Important here, these *super attractors* are independent of the 60 minutes data sets to be examined. Therefore, with the exception of the first minutes being influenced by the transient effect, the comparison should display results not much variating. And finally, the δM can be approximated by

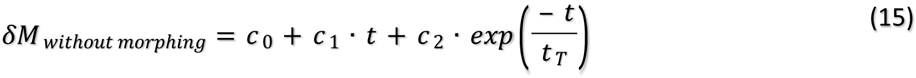

As before, the constants are approximated applying the Levenberg-Marquardt algorithm through the software CurveExpert Professional 2.6.5.

## Results

All input, measured data and simulation results, have a sampling frequency of 500 Hz. Further procedures, including generating graphs, where done after filtering with a ‘triple F low pass filter’ [27] with a cutoff frequency of 10 Hz.

A graphic comparison between measurement and simulation gives a fist impression of the model’s power (Fig 2).

**Fig 2.**
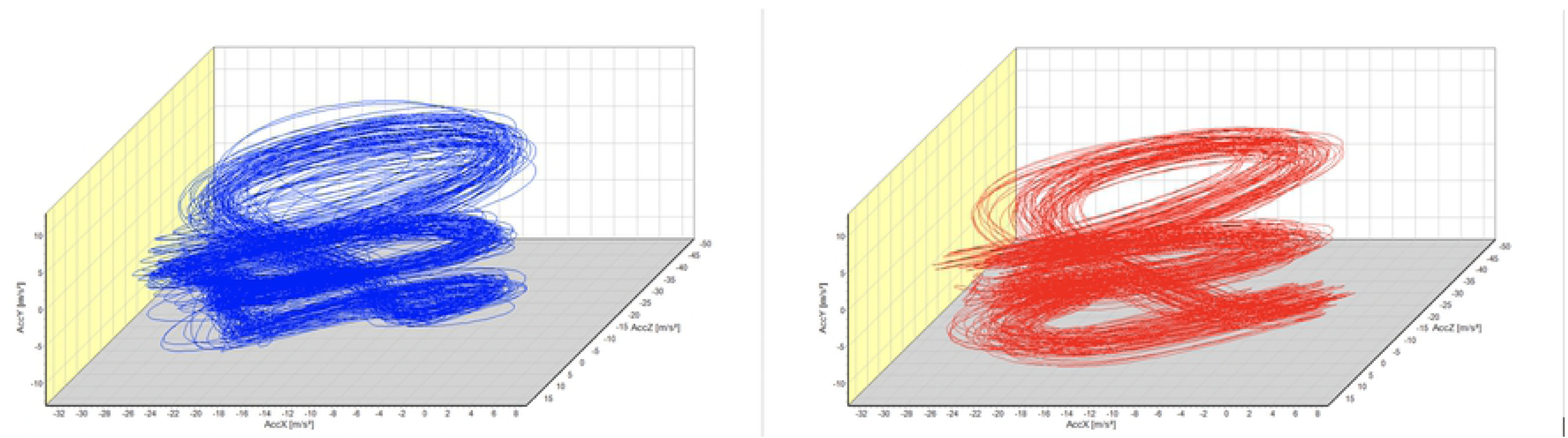
Measured data (blue) and simulation results (red) of the first run of subject three. The similarity analysis for running yield a gap between 50 and 56 % separating same and different subject comparisons (Fig 3). All comparisons, of measurements or simulations, between same subjects lie above the gap, comparisons between different subjects lie below.

**Fig 3.**
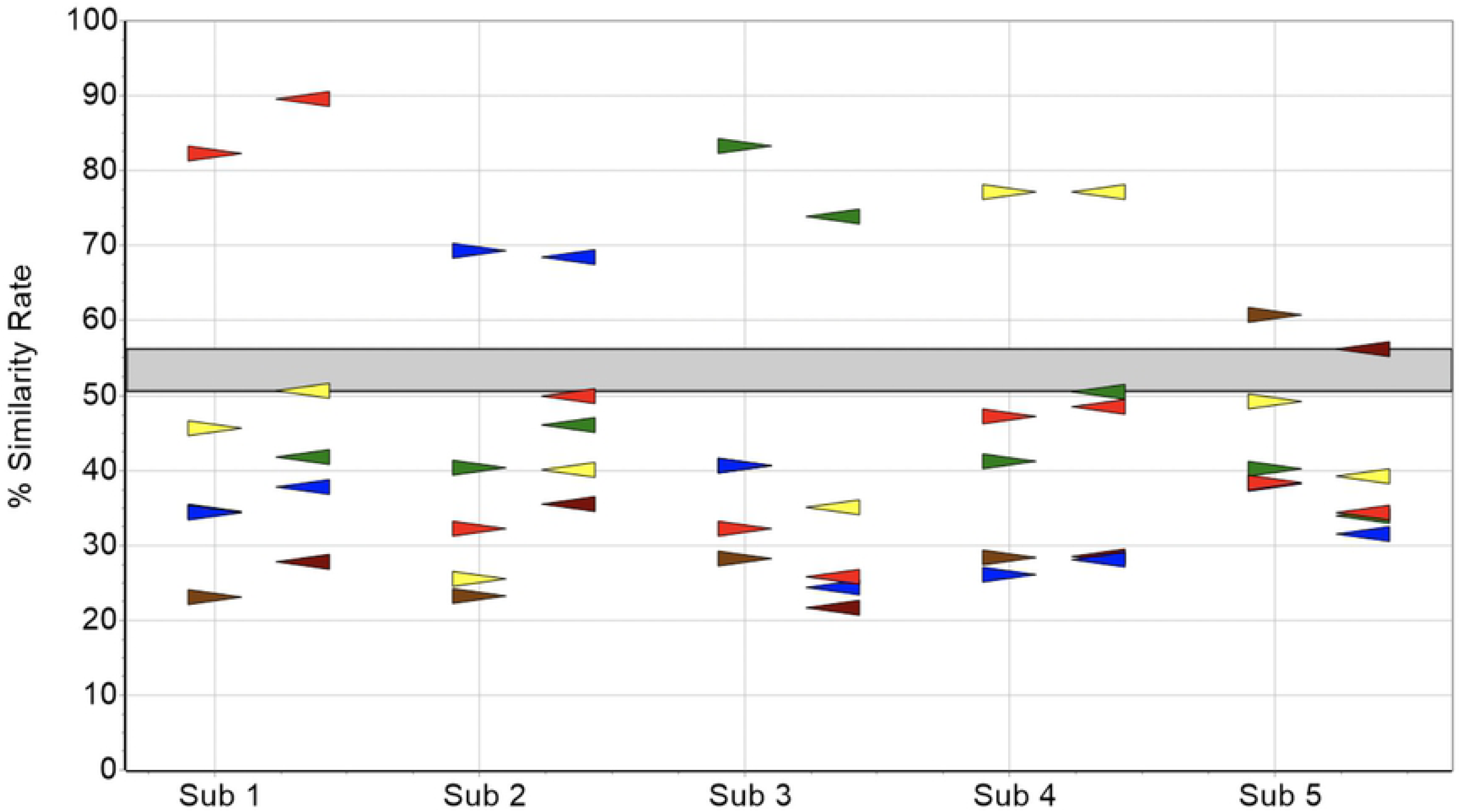
Similarity rate of running measurements (triangle right pointing) and simulations (triangle left pointing). For biking the seperating gap between same and different subject comparisons is 52 to 64 (Fig 4). As before all same subject comparisons lie above the gap, different subject comparisons below.

**Fig 4.**
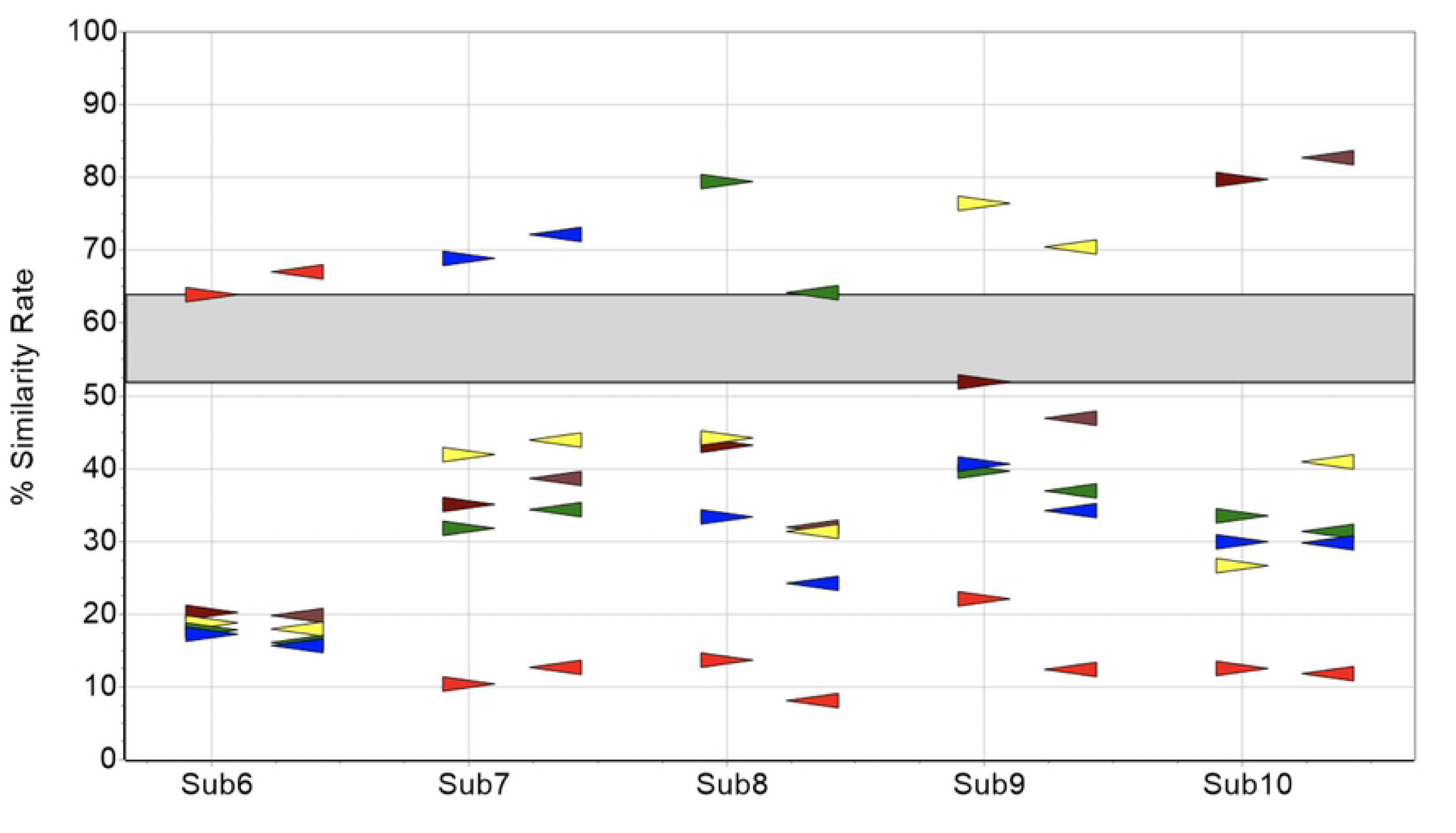
Similarity rate of biking measurements (triangle right pointing) and simulations (triangle left pointing). Values of *δM* – equation (14) – are influenced by morphing and transient effect. A typical progression with both factors influencing *δM*(*t*) is shown in Fig 5. In the first few minutes the transient effect causes an increase/decrease, while morphing with its more moderate decline is visible afterwards.

**Fig 5.**
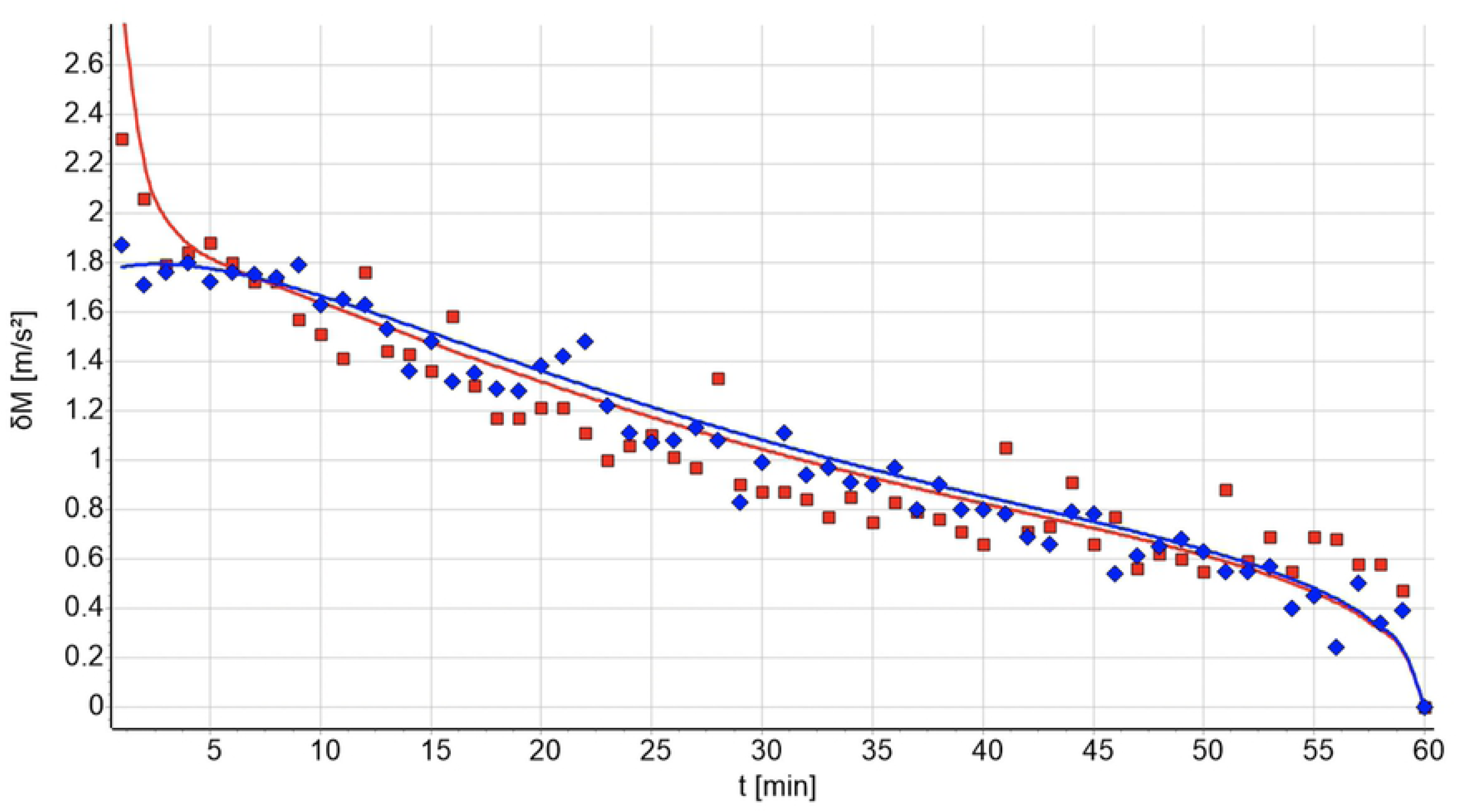
*δM*(*t*) for the measurement (blue) of one run and the respective simulation (red). The difference between measurement and the simulation are caused by the transient effect and the “short time fluctuation”. Here the starting conditions are largely random, causing differences at the beginning. The morphing of a specific measurement is imprinted into the simulation values via the equation (3). A morphing effect is visible, if the analyzed minutes are from one uninterrupted measurement. The comparisons with the *super attractor* calculated form data independent of the actual numbers do display random changes and the transient effect but no morphing (Fig 6). Those data can be approximated using equation (15), which allows to calculate the transient effect largely without the influence of morphing. δM does not variate much with the only remarkable deviation at the beginning up to about the 10^th^ minute.

**Fig 6.**
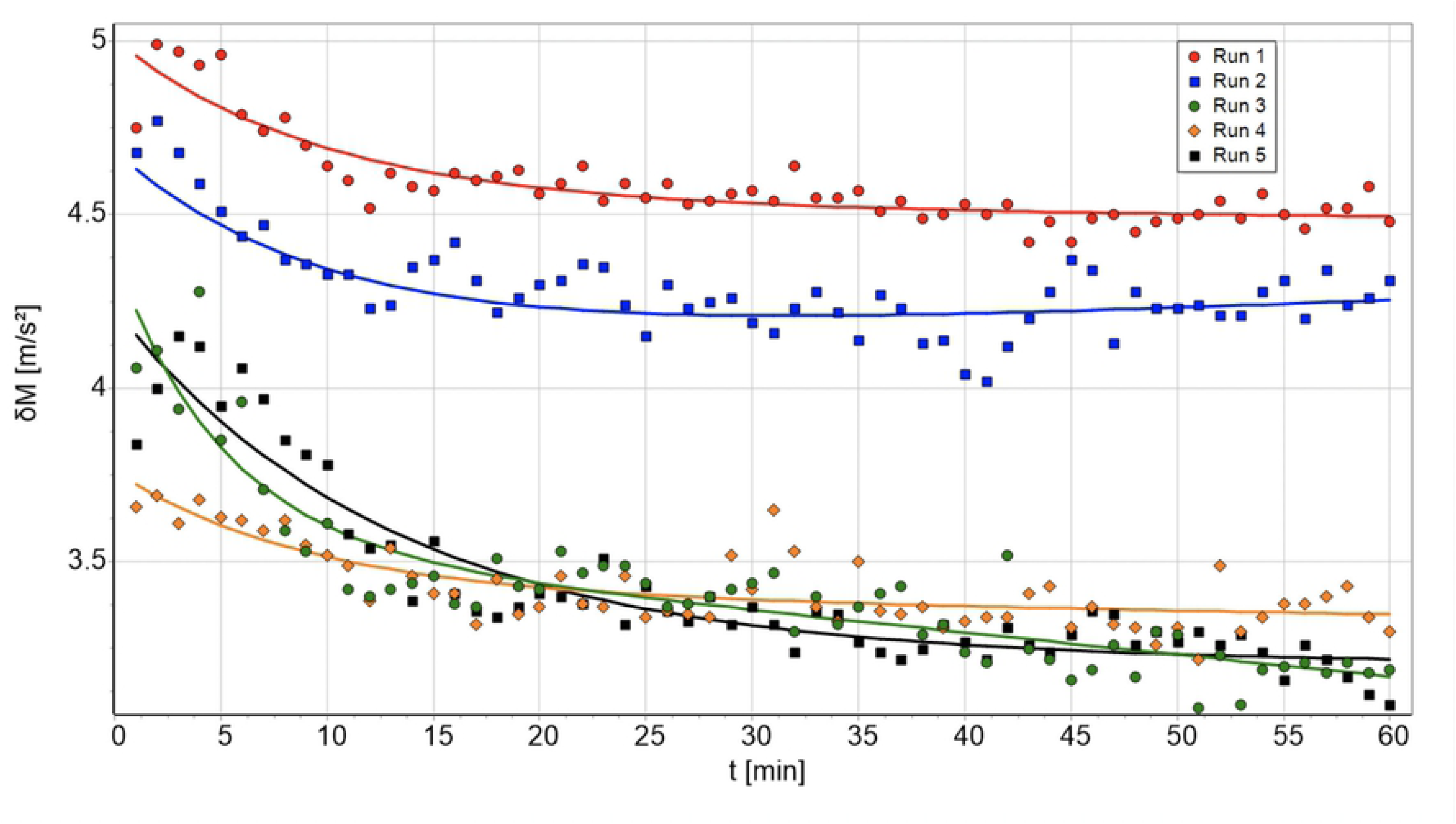
Five runs of subject 3 compared to the subject’s independent *super attractor*. The lines represent the approximation as of equation (15).Fig 6 shows the five values of subject 3 running, in which case a substantial transient effect is prominently visible. Other subjects show fewer or no exponential behavior at the beginning (Table 3).

**Table 3.** The time *t* _*T*_ [min] by which the transient effect reduces to *e* ^− 1^ of its start value.

| Sub | Run 1 | Run 2 | Run 3 | Run 4 | Run 5 | Sub | Bike 1 | Bike 2 | Bike 3 | Bike 4 | Bike 5 |
| --- | --- | --- | --- | --- | --- | --- | --- | --- | --- | --- | --- |
| Sub 1 | 1.0 | 5.1 | 3.9 | - | - | Sub 6 | - | - | - | - | - |
| Sub 2 | 5.0 | - | - | 5.0 | - | Sub 7 | - | - | - | - | - |
| Sub 3 | 9.7 | 10.0 | 5.1 | 9.0 | 12.9 | Sub 8 | 3.4 | 4.3 | 3.4 | 2.8 | 3.5 |
| Sub 4 | 1.7 | 1.8 | 0.9 | 5.5 | 1.0 | Sub 9 | - | - | - | - | - |
| Sub 5 | 1.9 | 2.8 | 3.7 | 2.3 | 3.5 | Sub 10 | - | 8.2 | 1.5 | 1.4 | - |

The absolute height of δM depends on the attractor’s similarity compared with the independent *super attractor*. The following graph (Figs 7 and 8) shows the average values of all subjects’ mean and the standard deviation for minutes 11 to 60 for all runs/bike trials.

**Fig 7.**
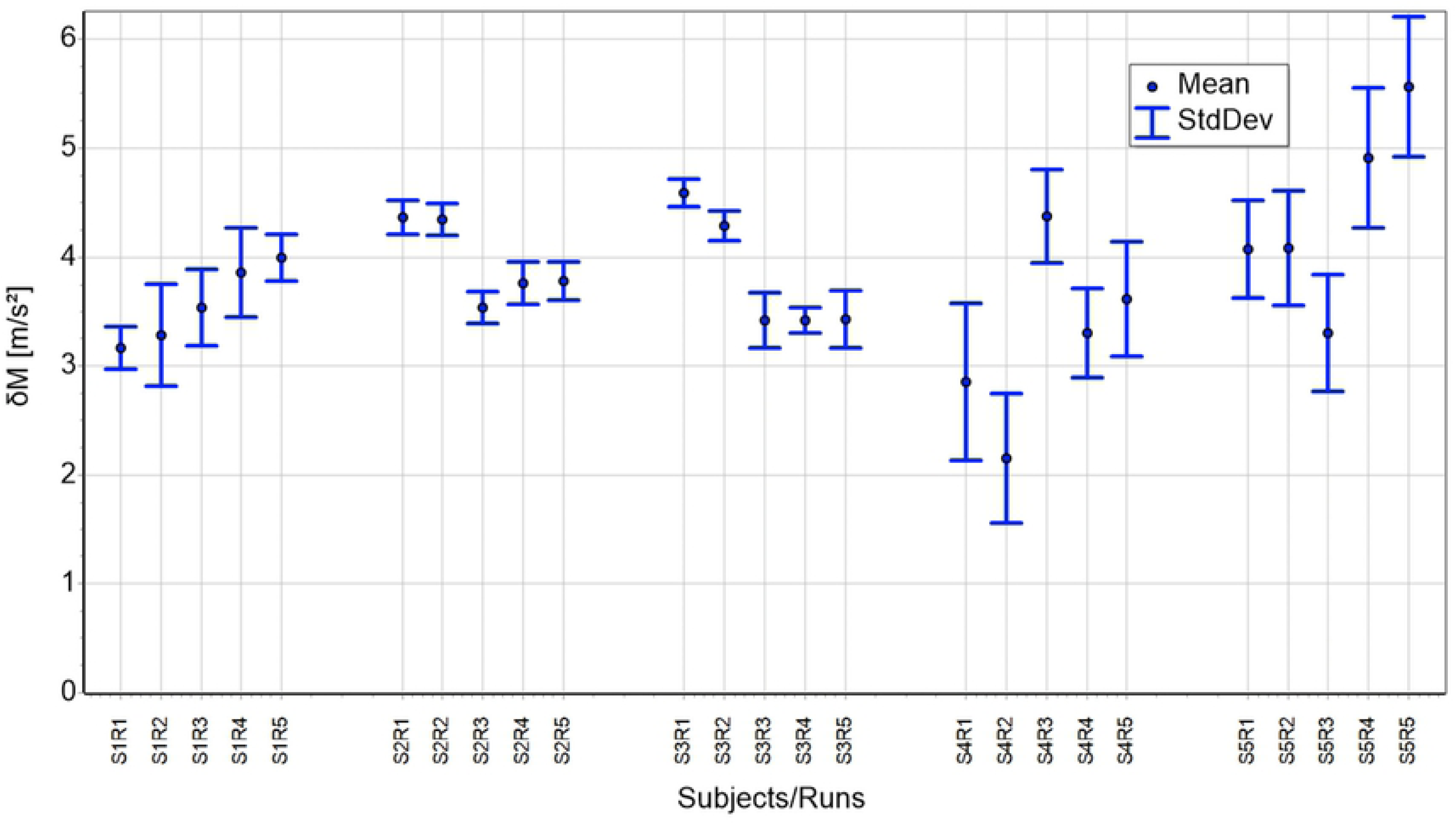
All 5 runs of all 5 subjects compared to their personal but independent super attractor for minutes 11 to 60.

**Fig 8.**
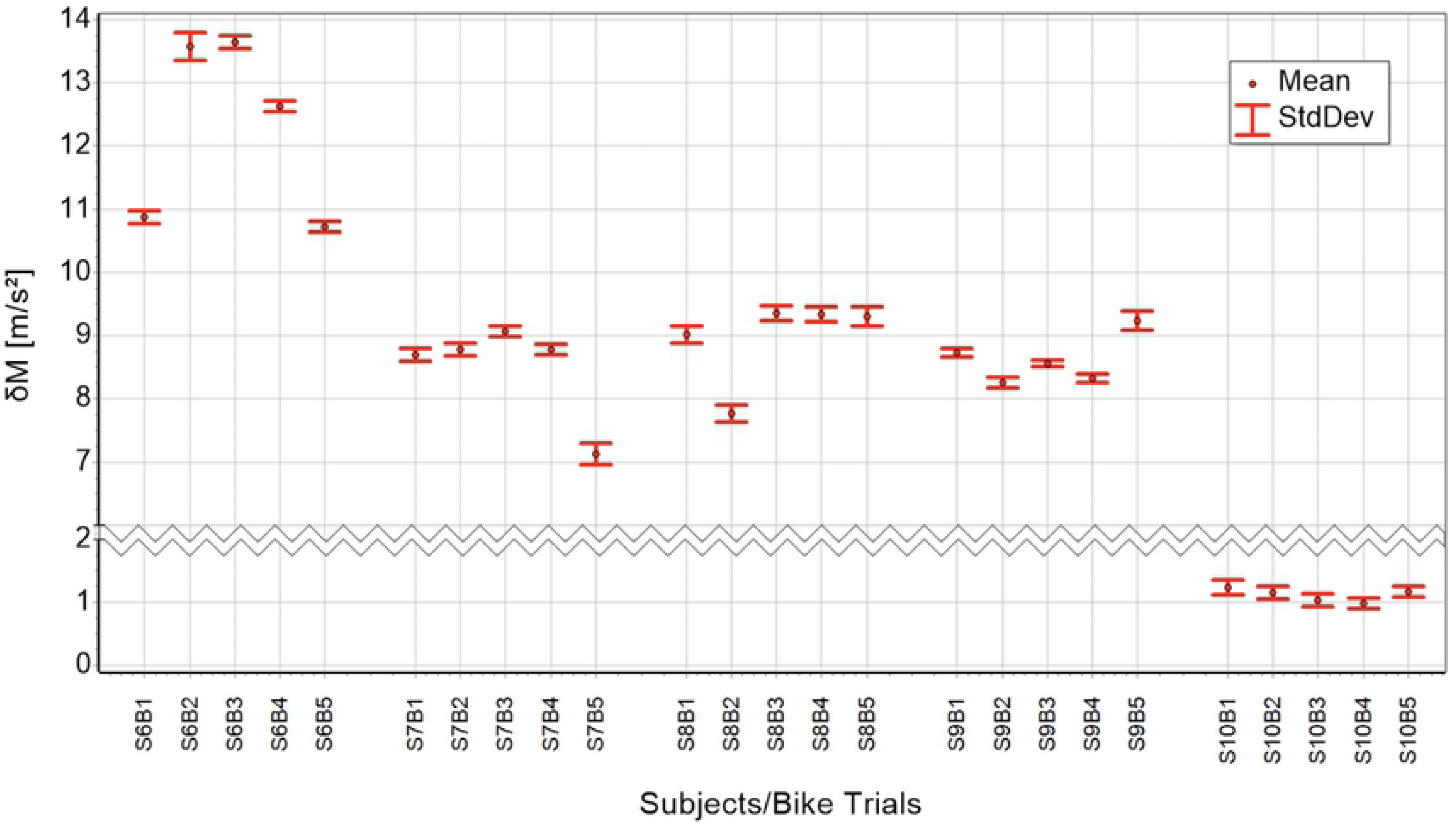
All 5 bike trials of all 5 subjects compared to their personal but independent super attractor for minutes 11 to 60.

## Discussion

The purpose of this paper is finding a quantitative description of cyclic motion with the capacity to simulate individuals’ characteristic movement. A model was proposed consisting of 6 contributing parts. Individual attractor, morphing, short time fluctuation, transient effect, control mechanism and sensor noise. Simulations based on this model showed the same distinctive variations as the measured data, confirming the model suitability for describing cyclic motion. The nine constants together with the subject’s attractor approximations are characteristic for a person’s movement and the influence of the recording sensors. As known from previous studies [21, 22] the influence of morphing and transient effect is small compared with the differences between individuals. While morphing is present in all trials, the transient effect is not observable in all cases. For biking the transient effect is less prominent compared to running. We suspect the fixation of the legs with the foot connected to the pedal and the hip very much fixed onto the saddle, there is not much freedom in movement variation. The tibia position and its associated acceleration is often settled onto the attractor from the start onwards. A different situation is seen in running, where the kinematic chain is unfixed near the location of the accelerometer at the distal end of the tibia. Here the probability to start a movement close to the subject’s attractor, resulting in no visible transient effect, is small. Interestingly, the most experienced runners show the least transient effect. The comparison of a subject’s attractors of a 1-hour measurement with an independent *super attractor* allows approximating the magnitude of morphing. The maximal difference between attractors from independent measurements of one subject is restricted by the maximal possible morphing. Morphing can deform an attractor in many different ways, which most probable result in *δM*’s of comparable values. Therefore, results as shown in Figs 7 and 8 might represent good approximations of medium morphing magnitudes. Still, the determination of the attractor remains a challenging issue. In mathematical systems, like the famous “Lorenz map”, the attractor is reached after the transient effect subsided. There, either a stable regular attractor is reached or a strange one is to be seen. Here, data of the cyclic motion never reaching compete regularity, neither is the behavior completely chaotic. The regularity is, as mentioned before, good enough to discriminate between individuals. Still the question remains, how to rate the attractors’ differences, when attractor approximations are calculated by averaging the cycles of different time intervals. Does it simply mean when doing the averaging over longer time periods these differences will almost completely vanish? Or, does it mean that attractors are changing with time, even if these changes are small? So far, we do not have enough data to answer this question with certainty. But, from the results above we rate the second statement as the more probable one. There is a theoretical argument for this statement as well. While developing the mathematical description of cyclic motion, the first approach was without morphing. The idea was having an attractor not depending on time and the fluctuation based on a “random walk” characteristic. This construct, however, did not allow for describing the full data variability. Altogether our model is capable describing cyclic motion quantitatively. Given the individual’s attractor approximations and the subject specific constants, simulations resulting in output, which is specific for the subject’s particular movement. In addition, there are other aspects needing further attention. One is establishing a threshold for the similarity analysis to define the percentage when recognition is achieved. Many more measurements of a specific cyclic movement should allow finding a suitable number by using the median method described by Vieten et al. [23]. Another limitation of the current approach is the focus on calculating δM, which depends on 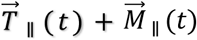, the parallel components only. Analyzing the full expression 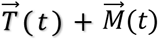 might allow further inside.

## Acknowledgment

We thank all subjects who participated in the study. The study was approved by the local Ethical Committee of the University of Konstanz, Germany under the RefNo: IRB19KN10-005.

## Author contribution

Developed the mathematical procedure: MMV. Conceived and designed the experiments: MMV, CW. Performed the experiments: CW. Analyzed the Data: MMV, CW. Wrote the paper: MMV, CW. Designed and programmed the software used in the analysis: MMV, CW.

